# Theta and alpha EEG oscillations reflect sleep need — except during the wake maintenance zone

**DOI:** 10.1101/2023.02.03.526951

**Authors:** Sophia Snipes, Elias Meier, Sarah Meissner, Hans-Peter Landolt, Reto Huber

## Abstract

Increasing time spent awake results in accumulated sleep need, a process known as sleep homeostasis. Sleep homeostasis combines with a 24 h circadian rhythm to determine when and for how long we sleep. Both sleep homeostasis and the circadian rhythm substantially affect spectral power of the wake electroencephalogram (EEG), but not in ways predicted by current models. Specifically, these models hypothesize that time spent awake increases neuronal synaptic strength, which increases synchronization and should therefore increase oscillatory activity. However, the dominant wake EEG oscillations, measured as theta (4-8 Hz) and alpha power (8-12 Hz), do not follow the predicted buildup in homeostatic sleep pressure with time awake. This is due to a limitation of spectral power analysis, which does not distinguish between changes in the amplitude of oscillations from changes in the quantity of oscillations present in the signal. We wished to determine whether the amplitudes of EEG oscillations would specifically reflect homeostatic sleep pressure, independently from changes in quantity. We collected data from 18 young healthy adults during a 4-h sleep / 24-h extended wake paradigm. We indeed found that theta and alpha oscillation amplitudes reflect homeostatic sleep pressure, increasing along a saturating exponential function with time awake. Instead, theta quantities increased linearly with time awake, and alpha quantities decreased. Notably, theta and alpha amplitudes temporarily decreased during the wake maintenance zone (WMZ), a 3-4 h time window just before bedtime when it is difficult to fall asleep. Using pupillometry, we also found that mean pupil diameter increased during this window, while variance decreased. These results suggest that the WMZ is dependent on an alerting signal from the ascending arousal system. The WMZ therefore counteracts the observed build-up in homeostatic sleep pressure reflected in EEG amplitudes by temporarily desynchronizing cortical activity.

## INTRODUCTION

Good sleep is essential for daily functioning and overall quality of life. The reason we need sleep is so that physiological systems used during the day have a dedicated period to rest and conduct structural maintenance (Vyazovskiy & Harris, 2013), clear metabolic by-products (Hauglund et al., 2020; Xie et al., 2013), restore overall functioning to baseline levels (Killgore, 2010; Van Dongen et al., 2003), and more. This is referred to as *sleep homeostasis* and is especially critical for the brain. Also important is the timing of sleep, controlled by a 24-h *circadian rhythm* which allow independent systems across the body and brain to synchronize their recovery in order to optimize overall performance. These homeostatic and circadian fluctuations make up the two-process model of sleep (Borbély, 1982). Together they create *sleep pressure*, which is the drive to fall asleep when needed and at the right time.

The homeostatic process in particular was designed to explain the notable changes in sleep slow waves, electroencephalographic (EEG) oscillations between 0.5 and 4 Hz that characterize NREM sleep (non rapid-eye-movement sleep, i.e. stages 2 & 3). Slow wave activity decreases exponentially during NREM sleep, reflecting homeostatic sleep pressure dissipation. Vice versa, slow wave activity at the beginning of sleep depends on the duration of prior wake, following an increasing saturating exponential function (Dijk et al., 1987). This means that the buildup in sleep need is steepest during the initial hours of wake, then gradually saturates with additional time awake (Figure 1A).

**Figure 1:**
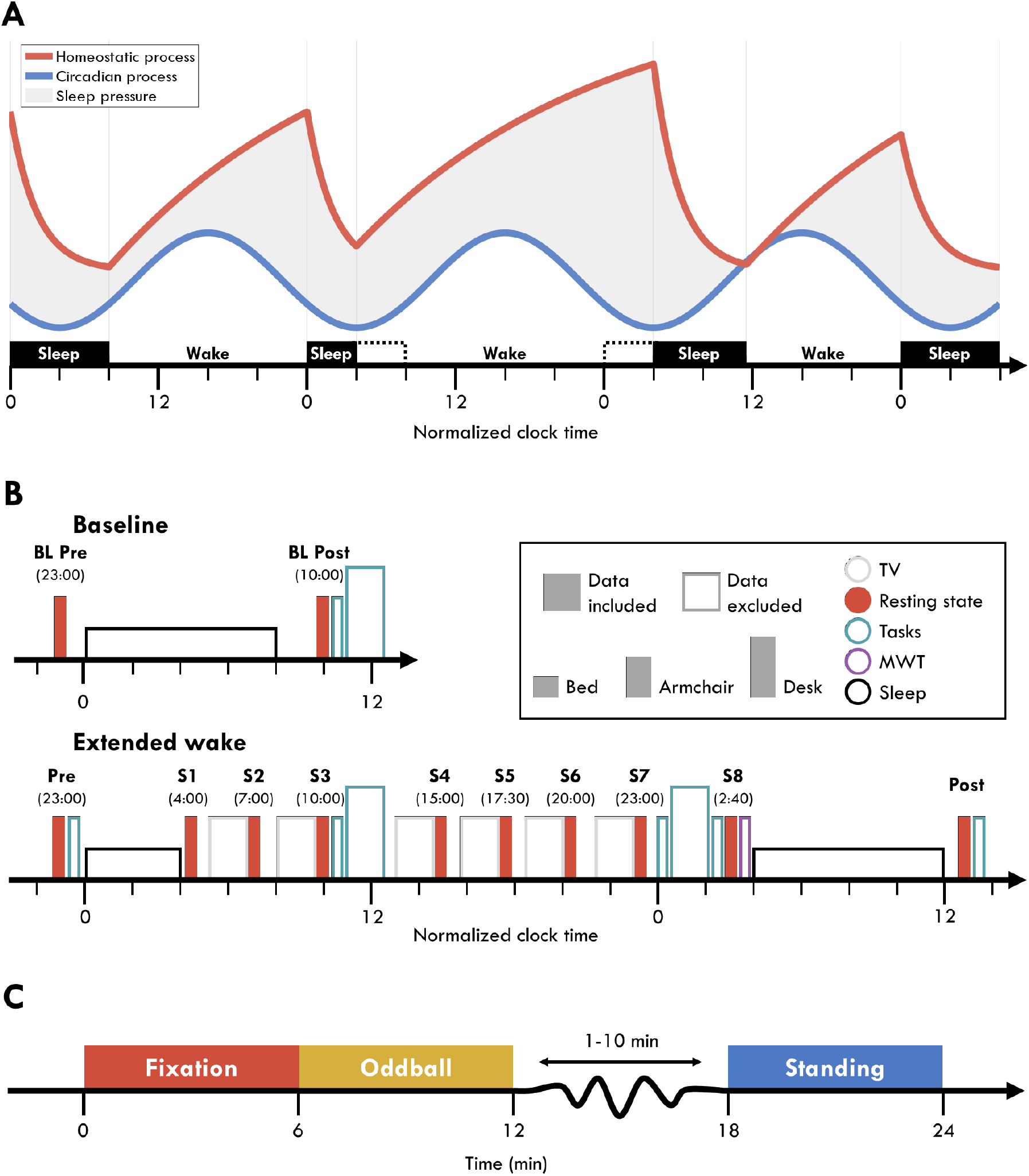
Experiment design. **A**) The two-process model during a 4/24 sleep/wake schedule. The red line reflects the homeostatic process, building sleep pressure monotonically with wake and dissipating during sleep. The blue line reflects the circadian process, peaking in the middle of the day and at its lowest in the middle of the night, independent of actual sleep and wake behavior. The shaded area reflects the resulting sleep pressure from combining these two processes. Black blocks indicate when participants actually slept, whereas the dotted outline indicates the window in which they would have slept according to their circadian rhythm. **B**) Experiment schedule. Each block indicates an EEG recording session. Filled blocks indicate data analyzed in this paper. Color indicates the activity participants engaged in: gray, watching TV; red, the resting state recordings in C; teal, task blocks analyzed in Snipes et al. (2022); purple, the MWT; black, sleep. The height of each block indicates the condition in which data was collected: short, lying in bed; medium, seated in a comfortable armchair with foot and backrest / standing; tall, seated at a desk. Brief empty spaces indicate transition periods allowing for delays. Six longer breaks were included prior to each TV block in which participants were provided with meals. Circadian time was normalized across participants to their habitual bedtime. Participants at baseline and during the recovery night were free to wake up when they wished, and at the beginning of the extended wake period they were woken up after 4 h of sleep. **C**) Timeline for the resting state recordings. Each condition was 6 minutes, and always done in the depicted order. Between the Oddball and Standing, a questionnaire was conducted which took a variable amount of time, followed by moving the participant from the armchair to standing. Abbreviations: BL, Baseline; MWT, Maintenance of Wakefulness Task.

A possible explanation for this increase in slow wave activity is that wakefulness progressively increases neuronal synaptic strength, which then requires sleep to restore overall synaptic balance. This is one of the key points of the synaptic homeostasis hypothesis (Tononi & Cirelli, 2003). In essence, learning during the day occurs through strengthening synaptic connections. Increased synaptic strength increases overall connectivity which leads to increased synchronicity of neuronal activity across the brain. This increased synchronicity between neurons will result in more synchronized oscillations in the surface EEG, detected as oscillations with larger amplitude with steeper slopes. Using computational models (Esser et al., 2007), animal sleep data (Vyazovskiy et al., 2007), and human sleep data (Riedner et al., 2007), the proponents of this hypothesis demonstrated how decreasing synaptic strength across sleep results in decreasing slow wave amplitudes and slopes. They argue that this buildup of synaptic connectivity during the day is one of the fundamental reasons sleep is needed: to restore neural synaptic homeostasis to baseline levels (Tononi & Cirelli, 2003, 2014).

While synaptic strength and sleep homeostasis can explain slow wave activity in sleep, they do not likewise explain changes in wake oscillations. Human wake EEG is predominantly characterized by alpha oscillations (8-12 Hz) and to a lesser extent theta oscillations (4-8 Hz), often measured as power in the frequency domain. Theoretically, the increased connectivity with time spent awake should affect these oscillations along a similar increasing saturating exponential function as for slow waves in sleep. However, while theta power does increase with sleep deprivation, the effect is rather linear (Finelli et al., 2000). Furthermore alpha power actually decreases (Cajochen et al., 2002).

In addition to neither oscillation following a homeostatic trajectory, both are also affected by circadian rhythmicity (Aeschbach et al., 1997; Cajochen et al., 2002; Strijkstra et al., 2003), further masking potential homeostatic effects. Alpha activity fluctuates in phase with core body temperature, a reliable circadian marker peaking in the middle of the day and lowest in the middle of the night (Åkerstedt et al., 1979; Cajochen et al., 2002). Instead, theta activity is lowest in the evening, corresponding to the wake maintenance zone (WMZ; Strogatz et al., 1987; Zeeuw et al., 2018). The WMZ, more dramatically known as the “forbidden sleep zone”, is a circadian window of 3-4 hours just prior to melatonin onset in which sleep becomes exceptionally difficult (Lavie, 1997). During the WMZ, sleep onset latencies substantially increase even during extensive sleep deprivation (Dijk & Czeisler, 1995; Lavie, 1986), subjective sleepiness decreases, and behavioral performance improves (McMahon et al., 2018, 2021; Shekleton et al., 2013; Zeeuw et al., 2018). The timing of the WMZ is not reflected in circadian markers such as melatonin levels or core body temperature, it has not been reported in any animal models to our knowledge, and is not represented in the classic two-process model of sleep.

Therefore, while increasing synaptic strength would have predicted an increase in both theta and alpha power with time spent awake, in practice neither oscillation strictly reflects this buildup in homeostatic pressure, and they are further synchronized to different circadian phases. However, the fact that wake oscillations do not reflect sleep homeostasis may be due to a limitation of spectral power analyses. “Power” refers to the amount of energy in a frequency band, and is typically calculated using some variant of the fast Fourier transform (M. X. Cohen, 2014). Once a time-series signal has been transformed into the frequency domain, power values are averaged or summed within a frequency range of interest, and this is the power for that band. While this is a simple and generally effective measure for quantifying oscillatory activity, it is simultaneously affected by the *quantity* of oscillations present in the signal and their *amplitude*, as well as broad-band changes in the entire spectrum (Donoghue et al., 2020).

The synaptic homeostasis hypothesis predicts that an increase in synaptic connectivity results in an increase in oscillatory amplitudes; this does not need to have any bearing on the number of oscillations that actually occur. It is therefore possible that non-homeostatic factors such as the WMZ could independently affect the quantity of oscillations, whereas time spent awake more specifically affects their amplitude. When both oscillation amplitudes and quantities change independently across wake recordings, the resulting power values will reflect some undifferentiated mix between the two. By separating these contributions, we may have a specific marker of homeostatic sleep pressure during wake. Not only would this provide supporting evidence for the hypothesis that sleep homeostasis is linked to synaptic plasticity, but also provide a marker for sleep pressure more easily acquired than slow wave activity during sleep.

We therefore wished to determine whether the circadian and homeostatic influences on theta and alpha oscillations could be dissociated in resting wake EEG by separately measuring changes in amplitude and changes in quantities of oscillations. Eighteen young healthy adults participated in a 4/24 extended wake paradigm (Figure 1A), in which they slept the first 4 hours of the night and were then kept awake for 24 hours with repeated resting state recordings (Figure 1B), while measuring high-density EEG. We conducted cycle-by-cycle analysis (Cole & Voytek, 2019) to identify bursts of oscillations in the theta and alpha range, a method which identifies oscillations based on the morphology of the EEG signal rather than relying on power and amplitude thresholds. We then looked at changes in the mean amplitude of bursts and the average number of cycles (i.e., oscillations present in a burst) per minute for each band. Our prediction was that both theta and alpha amplitudes would follow an increasing saturating exponential function across extended wake, and show decreases following sleep. At the same time, the decrease in alertness with time awake should result in a decrease in the overall number of alpha oscillations. Likewise, circadian changes such as the decrease in theta during the WMZ should be reflected in decreases in the number of bursts.

To independently monitor changes in alertness across the extended wake period, we also recorded pupillometry with infrared cameras. Pupillometry and ocular behavior are indirect but rich signals thought to reflect activity of deep-brain nuclei. Pupil diameter and pupil responses to salient stimuli have been linked to alertness-promoting activity in the locus coeruleus (Aston-Jones & Cohen, 2005; Joshi et al., 2016; Murphy et al., 2014), as well as other interconnected nuclei in the brainstem and forebrain that make up the ascending arousal system (AAS) (Lloyd et al., 2022; Reimer et al., 2016). Blink rates have been shown to be an indirect measure of dopamine function (Jongkees & Colzato, 2016), and were hypothesized to reflect a compensation mechanism to counteract sleep deprivation (Barbato et al., 2007). Similarly, extended eye-closure can reflect microsleeps (Ong et al., 2013), which can anticipate the onset of whole-brain sleep (Skorucak et al., 2020). While there are still many open questions about the link between ocular behavior and sleep/wake promoting nuclei, the relationship between any of these signals and EEG could help better explore the underlying mechanisms driving the changes in oscillatory activity across extended wake. In short, while the two-process model and the synaptic homeostasis hypothesis provide specific predictions about wake EEG amplitudes, with pupillometry we hoped to provide possible explanations for changes in oscillatory occurrences.

## RESULTS

We recorded EEG, pupillometry, and questionnaire data from participants performing 3 wake resting state recordings, for 6 minutes each (Figure 1C). The first was a standard recording, *Fixation*, in which participants were seated in a comfortable armchair and had to gaze at a fixation point ~3 m away. The second was an active auditory *Oddball*, in which tones were presented randomly and participants had to push a button after “oddball” (i.e. deviant) tones. Afterwards participants filled out a questionnaire, and then got up for the final recording of *Standing* with eyes closed. They were asked to stand for this condition because during sleep deprivation participants quickly fall asleep with eyes closed. These rest recordings were conducted 12 times: before and after each of the 3 nights of sleep, and 6 more times throughout the extended wake period, approximately 3 hours apart (Figure 1B). The primary focus of this study was the Fixation condition, used in previous studies. The other two conditions were included to establish to what extent the observed effects extended to different resting states.

Participants maintained a regular sleep-wake schedule the week prior to each experiment bout, and therefore the timing of the individual 24 h circadian sleep rhythm could be inferred from previous studies (Åkerstedt et al., 1979; Dijk & Czeisler, 1995; Wyatt et al., 1999). Changes synchronized to melatonin would be high during night recordings (S1, S2, S8), and low during day recordings. Vice versa, circadian changes synchronized to core body temperature would peak in the middle of the day (S4, S5), and be low in the middle of the night (S1, S8). Because the focus was on homeostatic changes, these comparisons were not statistically analyzed, but can be inferred from the outcome measures’ trajectories.

To quantify the effects of extended wake, sleep, and the WMZ, for each outcome measure we conducted the same three paired t-tests. For wake-dependent changes, we compared values from the start and end of the 24 h extended wake period, S1 and S8. These were within 2 h of the same circadian phase, therefore any differences should largely be due to sleep homeostasis. To quantify sleep-dependent changes, we compared values from the wake recordings before and after the baseline night, BL Pre and BL Post. Unlike for wake changes, these were conducted at different circadian times and the difference in sleep homeostatic pressure was lower, however these are typical recordings during sleep studies.

To statistically quantify any deviation during the WMZ from the underlying trajectory of a given outcome measure, we linearly interpolated values from S5 (17:30) to S8 (2:40) for timepoint 21:30 and compared it to the average of S6 (20:00) and S7 (23:00). The timepoints of the WMZ were determined based on the converging results of subjective sleepiness (Figure 2) and theta power (Figure 3A), both of which showed a decrease in an otherwise monotonic increase during recordings S6 and S7, corresponding to 1-4 h before habitual bedtime.

**Figure 2:**
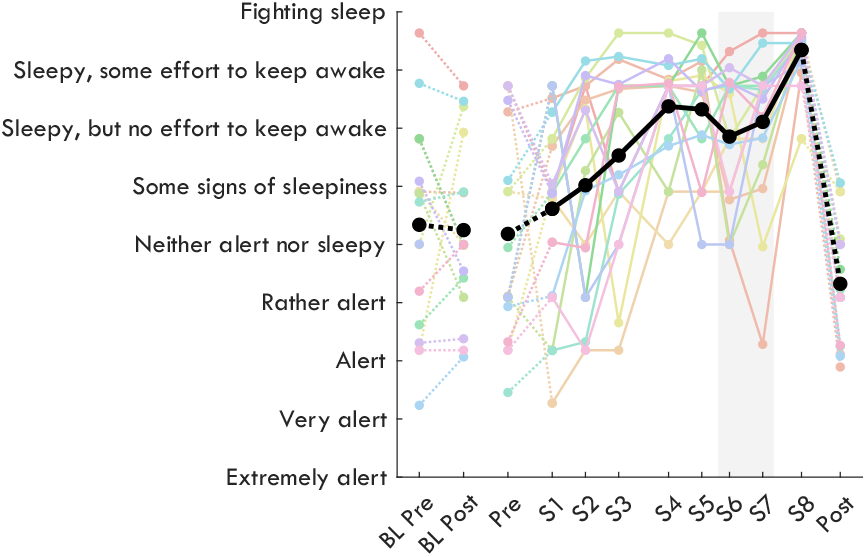
Subjective sleepiness. Sleepiness was measured on a continuous visual-analog adaptation of the KSS, using the original labels as markers (y-axis). The thick black line indicates the group average, and thin colored lines are datapoints of individual participants. Solid lines connect sessions during the same-day extended wake period, and dashed lines indicate changes across sleep. S1-S8 are spaced out relative to the time they occurred within the 24 h wake period (Figure 1B). The shaded gray area indicates the WMZ. Acronyms: BL, baseline; KSS, Karolinska Sleepiness Scale; WMZ, wake maintenance zone.

**Figure 3:**
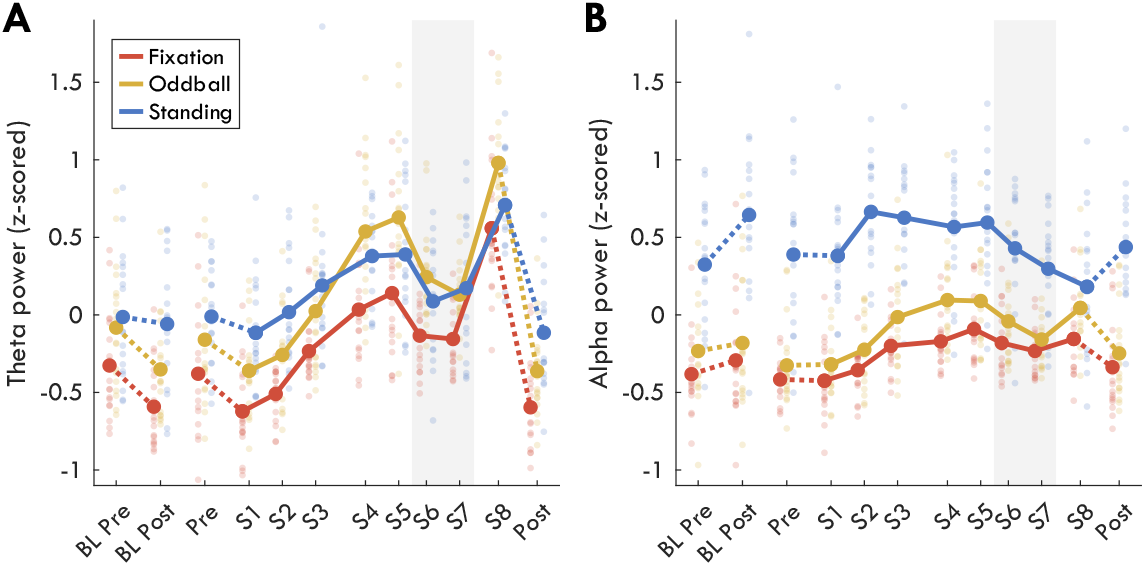
Z-scored power band changes. **A**) Theta power (4-8 Hz) and **B**) alpha power (8-12 Hz). Thick lines indicate group averages for each condition (as different colors) across sessions (x-axis). Solid lines connect sessions during the same-day extended wake period, and dashed lines indicate changes across sleep. S1-S8 are spaced out relative to the time they occurred within the 24 h wake period (Figure 1B). Dots reflect individual participants’ datapoints. The shaded gray area indicates the WMZ. Power spectral density values were first z-scored for each frequency pooling channels, sessions, and conditions. All channels were included in the average except edge channels: 48, 63, 68, 73, 81, 88, 94, 99, 119. Finally, z-scored values within each band range were averaged. Acronyms: BL, baseline; WMZ, wake maintenance zone.

All t-values, degrees of freedom, p-values, and Hedge’s g effect sizes are provided together in Table 1. Throughout the text, only the corresponding t-values will be reported, unless either effect sizes or p-values are specifically of interest (e.g. when trending).

**Table 1:** Statistics results. Paired t-tests were conducted to determine overnight changes at baseline (BL Pre vs BL Post), changes across 24 h of extended wake (S1 vs S8), and deviations from the wake trajectories during the WMZ (S5&S8 vs S6&S7). All values were z-scored for each participant, pooling sessions, and conditions. Power values were z-scored separately for each frequency prior to being averaged into bands, and pupil oddball responses were z-scored also across timepoints prior to measuring the average the response. Degrees of freedom are specified in the subscript of t-values and reflect the number of datapoints for each comparison (N = DF + 1). Effect sizes are provided as Hedge’s g values. All statistics are with α = 5%. There is no correction for multiple comparisons. Acronyms: WMZ, wake maintenance zone; BL, baseline; SD, standard deviation.

|  | Condi-<br>tion | Overnight baseline | Extended wake | WMZ |
| --- | --- | --- | --- | --- |
| **Sleepiness** | – | $t_{(15)} = -0.21, p = .833, g = -0.06$ | $t_{(16)} = 7.36, p < .001, g = 2.61$ | $t_{(16)} = -3.36, p = .004, g = -1.13$ |
| **Theta power** | Fixation | $t_{(17)} = -4.80, p < .001, g = -0.83$ | $t_{(16)} = 9.66, p < .001, g = 2.99$ | $t_{(16)} = -5.96, p < .001, g = -1.78$ |
| | Oddball | $t_{(17)} = -3.19, p = .005, g = -0.70$ | $t_{(16)} = 10.41, p < .001, g = 3.76$ | $t_{(16)} = -6.92, p < .001, g = -2.17$ |
| | Standing | $t_{(16)} = -0.57, p = .574, g = -0.14$ | $t_{(16)} = 5.97, p < .001, g = 2.26$ | $t_{(16)} = -5.19, p < .001, g = -1.21$ |
| **Alpha power** | Fixation | $t_{(17)} = 1.75, p = .098, g = 0.26$ | $t_{(16)} = 3.03, p = .008, g = 1.07$ | $t_{(16)} = -2.86, p = .011, g = -0.63$ |
| | Oddball | $t_{(17)} = 0.82, p = .421, g = 0.13$ | $t_{(16)} = 3.89, p = .001, g = 1.32$ | $t_{(16)} = -3.61, p = .002, g = -0.89$ |
| | Standing | $t_{(16)} = 2.57, p = .020, g = 0.71$ | $t_{(16)} = -1.44, p = .168, g = -0.49$ | $t_{(16)} = -1.13, p = .276, g = -0.16$ |
| **Theta burst am-**<br>**plitude** | Fixation | $t_{(15)} = -2.06, p = .057, g = -0.74$ | $t_{(16)} = 6.71, p < .001, g = 2.40$ | $t_{(16)} = -3.19, p = .006, g = -0.70$ |
| | Oddball | $t_{(16)} = -1.99, p = .064, g = -0.47$ | $t_{(15)} = 6.92, p < .001, g = 2.51$ | $t_{(16)} = -4.78, p < .001, g = -1.50$ |
| | Standing | $t_{(16)} = -2.55, p = .022, g = -0.55$ | $t_{(16)} = 2.15, p = .047, g = 0.80$ | $t_{(16)} = -0.46, p = .655, g = -0.10$ |
| **Alpha burst**<br>**amplitude** | Fixation | $t_{(17)} = -5.55, p < .001, g = -0.75$ | $t_{(16)} = 4.49, p < .001, g = 1.65$ | $t_{(16)} = -3.71, p = .002, g = -0.97$ |
| | Oddball | $t_{(17)} = -2.00, p = .061, g = -0.43$ | $t_{(16)} = 7.52, p < .001, g = 2.18$ | $t_{(16)} = -6.47, p < .001, g = -1.58$ |
| | Standing | $t_{(16)} = 0.14, p = .889, g = 0.04$ | $t_{(16)} = 0.17, p = .870, g = 0.06$ | $t_{(16)} = -2.22, p = .041, g = -0.32$ |
| **Theta burst cy-**<br>**cles/min** | Fixation | $t_{(17)} = 1.08, p = .295, g = 0.34$ | $t_{(16)} = 5.57, p < .001, g = 2.00$ | $t_{(16)} = -1.77, p = .095, g = -0.58$ |
| | Oddball | $t_{(17)} = 1.25, p = .227, g = 0.30$ | $t_{(16)} = 7.80, p < .001, g = 2.52$ | $t_{(16)} = -4.03, p = .001, g = -1.24$ |
| | Standing | $t_{(16)} = 0.67, p = .513, g = 0.24$ | $t_{(16)} = 4.68, p < .001, g = 1.70$ | $t_{(16)} = -5.02, p < .001, g = -1.66$ |
| **Alpha burst cy-**<br>**cles/min** | Fixation | $t_{(17)} = 3.32, p = .004, g = 0.96$ | $t_{(16)} = -2.87, p = .011, g = -1.13$ | $t_{(16)} = 1.98, p = .065, g = 0.67$ |
| | Oddball | $t_{(17)} = 2.63, p = .017, g = 0.77$ | $t_{(16)} = -3.31, p = .004, g = -1.37$ | $t_{(16)} = 2.51, p = .023, g = 0.48$ |
| | Standing | $t_{(16)} = 3.63, p = .002, g = 0.89$ | $t_{(16)} = -6.17, p < .001, g = -1.73$ | $t_{(16)} = 2.91, p = .010, g = 0.56$ |
| **Pupil diameter**<br>**(mean)** | Fixation | $t_{(13)} = 0.39, p = .699, g = 0.13$ | $t_{(16)} = -3.78, p = .002, g = -1.22$ | $t_{(15)} = 4.65, p < .001, g = 1.26$ |
| | Oddball | $t_{(12)} = -1.79, p = .099, g = -0.47$ | $t_{(12)} = -1.39, p = .189, g = -0.59$ | $t_{(12)} = 2.17, p = .051, g = 0.64$ |
| **Pupil diameter**<br>**(SD)** | Fixation | $t_{(13)} = -3.76, p = .002, g = -0.98$ | $t_{(16)} = 3.52, p = .003, g = 1.25$ | $t_{(15)} = -1.69, p = .111, g = -0.65$ |
| | Oddball | $t_{(12)} = -3.66, p = .003, g = -1.22$ | $t_{(12)} = 4.74, p < .001, g = 1.56$ | $t_{(12)} = -2.58, p = .024, g = -0.90$ |
| **Pupil oddball**<br>**response** | Oddball | $t_{(9)} = 0.26, p = .802, g = 0.07$ | $t_{(11)} = -1.58, p = .143, g = -0.55$ | $t_{(9)} = 2.08, p = .067, g = 0.74$ |
| **Blink rate** | Fixation | $t_{(13)} = -1.32, p = .209, g = -0.39$ | $t_{(16)} = 0.62, p = .545, g = 0.22$ | $t_{(15)} = -1.25, p = .229, g = -0.42$ |
| | Oddball | $t_{(13)} = -0.02, p = .987, g = -0.01$ | $t_{(13)} = 4.37, p = .001, g = 1.44$ | $t_{(15)} = -0.21, p = .839, g = -0.08$ |
| **Ocular micro-**<br>**sleeps (%)** | Fixation | $t_{(13)} = -0.97, p = .350, g = -0.32$ | $t_{(16)} = 4.81, p < .001, g = 1.56$ | $t_{(15)} = -4.85, p < .001, g = -1.68$ |
| | Oddball | $t_{(13)} = -1.32, p = .210, g = -0.47$ | $t_{(13)} = 4.18, p = .001, g = 1.41$ | $t_{(15)} = -6.28, p < .001, g = -2.16$ |

### Changes in theta power but not alpha power replicate previous results

Before investigating oscillatory burst activity, we first determined whether our novel experimental paradigm replicated findings of previous studies showing both circadian and homeostatic changes in theta and alpha power. We expected an increase in theta and a decrease in alpha with increasing time awake, as well as a dip in theta during the WMZ, and a peak in alpha in the middle of the day (Cajochen et al., 2002). Power spectral density was calculated using Welch’s method for every channel during each recording. These values were z-scored separately for each frequency, pooling channels, conditions, and sessions. Z-scored power values were then averaged across channels, and then averaged within the theta and alpha bands.

Changes in theta power are plotted in Figure 3A. After the baseline night, there was a significant decrease in theta power for the Fixation (t_(17)_ = −4.80) and Oddball recordings (t_(17)_ = −3.19), but no change during Standing with eyes closed (t_(16)_ = −0.57). Across extended wake there was a substantial increase in theta power in all conditions (Fixation, t_(16)_ = 9.66; Oddball, t_(16)_ = 10.41; Standing, t_(16)_ = 5.97). During the WMZ, all conditions showed very large and significant decreases in theta power (Fixation, t_(16)_ = −5.96; Oddball, t_(16)_ = −6.92; Standing, t_(16)_ = −5.19).

Changes in alpha power are plotted in Figure 3B. After the baseline night, alpha power showed a trend increase for Fixation (t_(17)_ = 1.75, p = .098), no change for Oddball (t_(17)_ = 0.82), and a significant increase during Standing (t_(16)_ = 2.57). Across extended wake, alpha actually increased for Fixation (t_(16)_ = 3.03) and Oddball (t_(16)_ = 3.89) and showed no significant change during Standing, although on average decreased (t_(16)_ = −1.44). A significant dip in alpha was present during the WMZ in the Fixation condition (t_(16)_ = −2.86) and even more prominent in the Oddball (t_(16)_ = −3.61).

Given the discrepancy with previous results that found decreases in alpha with sleep deprivation (Cajochen et al., 2002; Strijkstra et al., 2003), we inspected the spectrograms of z-scored power to determine whether some other factor was contributing to the increase in alpha power in our data. The power spectrums for each condition and for three regions of interest (ROI; Front, Center, and Back) across the 8 extended wake sessions are plotted in Figure 4. Theta created a distinct peak in the spectrum in the Front ROI, and alpha created a prominent peak in the Back ROI, especially in the eyes-closed Standing condition. During S8, both a theta and an alpha peak was present in the Back ROI. Notably, the alpha peak did in fact decrease in amplitude following extended wake in the Fixation and Oddball Back ROI, but at the same time there were broad-spectrum increases from S1 to S8. Furthermore, the increase in both theta and beta (15-30 Hz) bled into the alpha range, especially in the Front and Center ROI. The change in spectrum was more unusual during the Standing recordings, with S8 not showing the same distinctive alpha peak in the Back ROI, but rather a new 6-10 Hz peak instead. These results confirm that theta and alpha were distinct oscillations. However, there seem to be also broadband increases in power with extended wake, and the changes in neighboring bands likely affected average alpha power as well.

**Figure 4:**
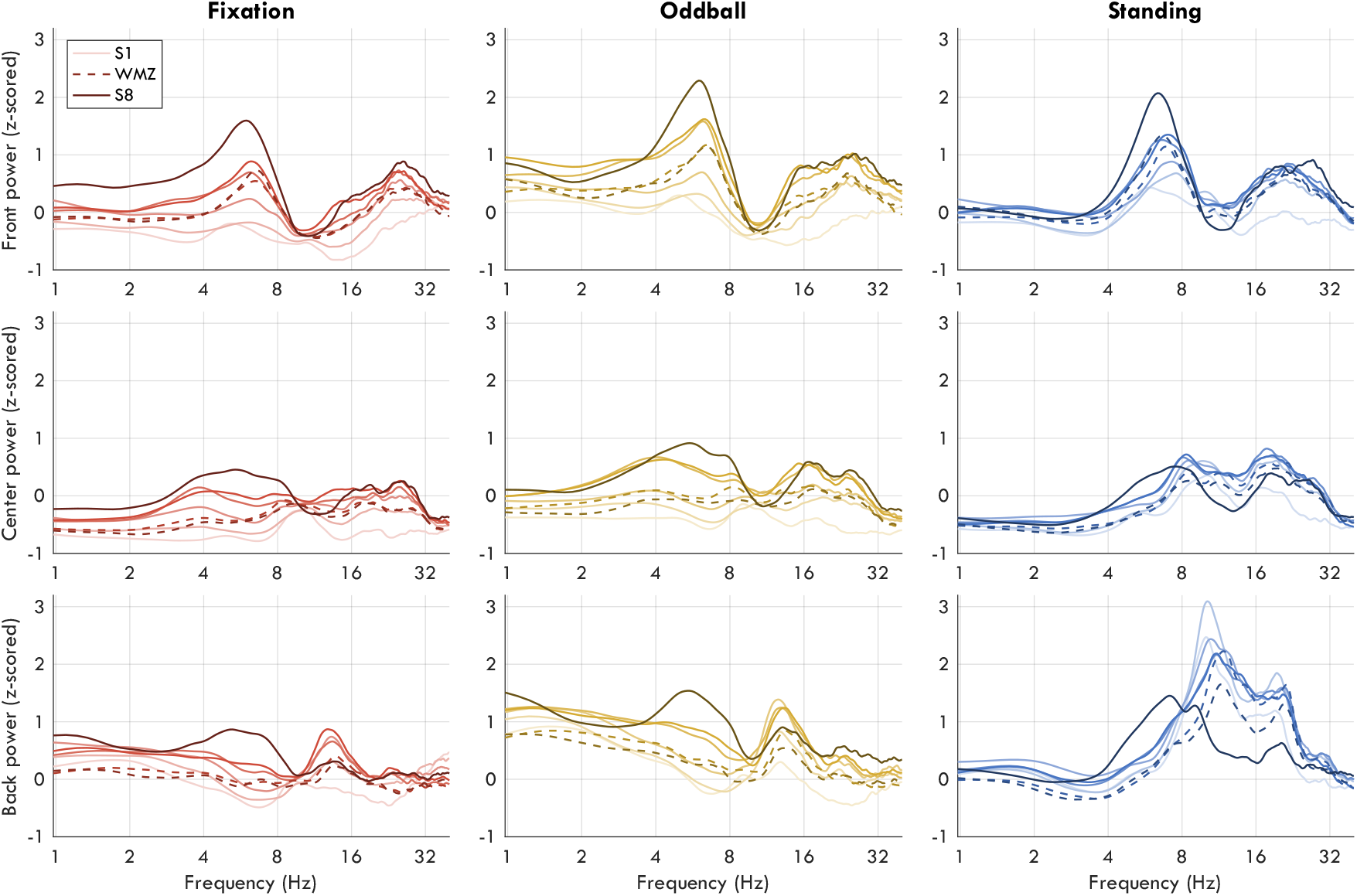
Z-scored power spectrums across extended wake. Each row plots an ROI (Front, Center, Back), each column a different condition. Color darkness indicates session, from S1 to S8, such that darker lines indicate more time awake. Dashed lines are the WMZ recordings (S6, S7). The x-axis indicates frequency on a log scale. Exact channels of the ROIs are provided in Snipes et al. (2022). Acronyms: WMZ, wake maintenance zone; ROI, region of interest.

### Cycle-by-cycle analysis successfully captures oscillatory activity

Cycle-by-cycle analysis was used to identify bursts between 2 and 14 Hz. Figure 5 provides an example of the EEG and burst detection during S8. Analyses were then performed on bursts between 4 and 8 Hz for theta, and 8 and 12 Hz for alpha. Oscillation amplitudes were quantified as the average negative-to-positive peak amplitude for all the cycles involved in a burst. The “number” of oscillations were quantified as the number of cycles per minute.

**Figure 5:**
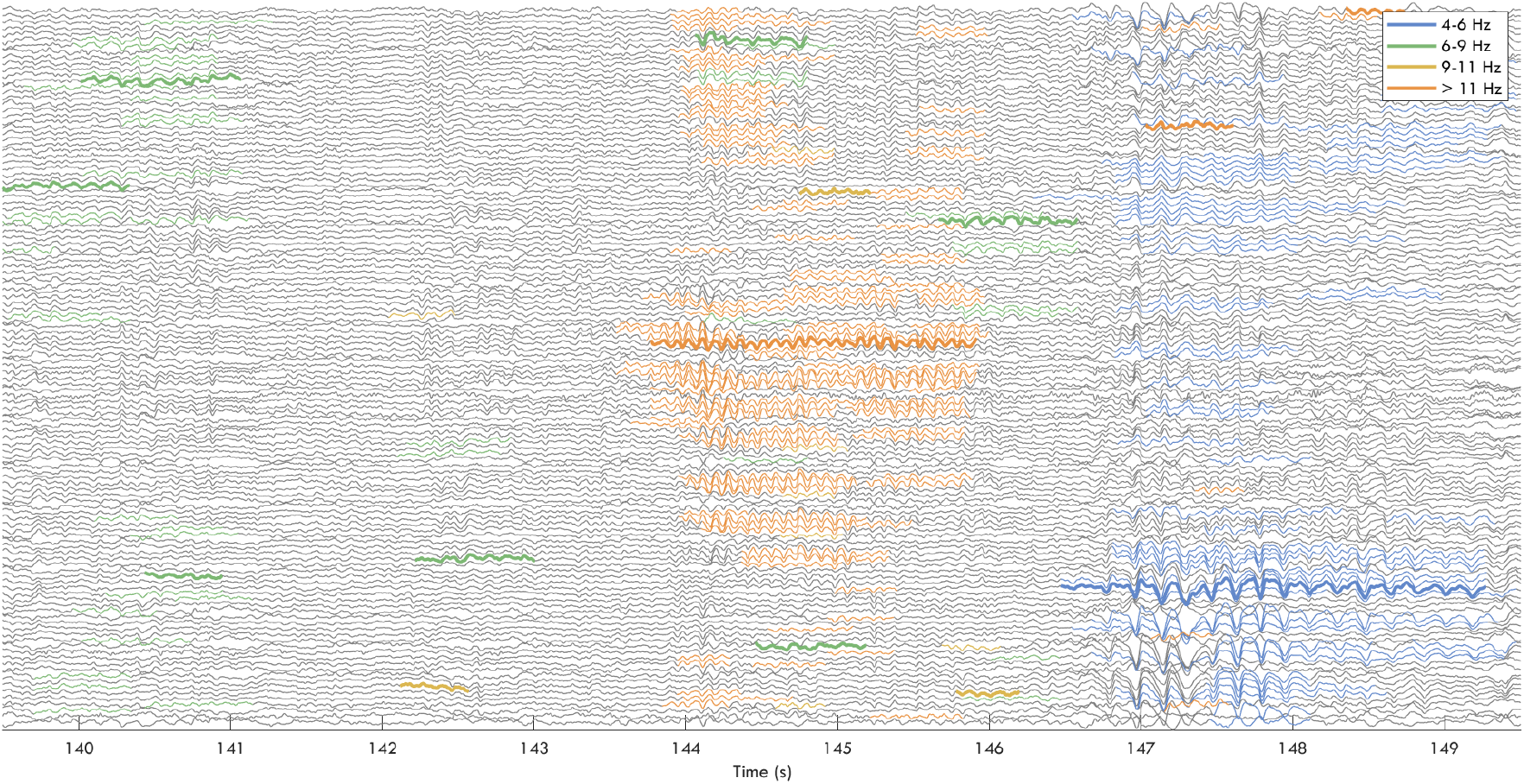
Example of detected bursts. 10 seconds of data from P15 Fixation (see Figure 6A). EEG data traces are in gray. Thick colored lines indicate the “reference” burst, the longest among temporally overlapping bursts in the same channel. Thin colored lines indicate overlapping bursts across channels considered to be the “same” burst as the reference. These were associated with the reference because mean frequencies were within 1 Hz of each other. Different colors represent different frequency bands. Acronyms: EEG, electroencephalography.

Figure 6A-B plots the distribution of the number of bursts by frequency for two example participants. Cycle-by-cycle analysis allowed clear differentiation between clusters of bursts by frequency. However, many individuals did not have two (theta/alpha), but in fact three or more clusters, and these distributions changed with time awake (best example: Figure 6A, Fixation). At the same time, other participants showed more classical bimodal distributions (Figure 6B). Ideally, we would have used individualized bands to delineate theta and alpha, however these shifting distributions complicate such an approach. Therefore, we limited ourselves to group average results using traditional frequency bands. Furthermore, rather than quantifying the occurrence of a given oscillation by the number of bursts as shown in Figure 6A-C, we used instead the number of cycles per minute (Figure 6G), as this also captures changes in burst duration (Figure 6E).

**Figure 6:**
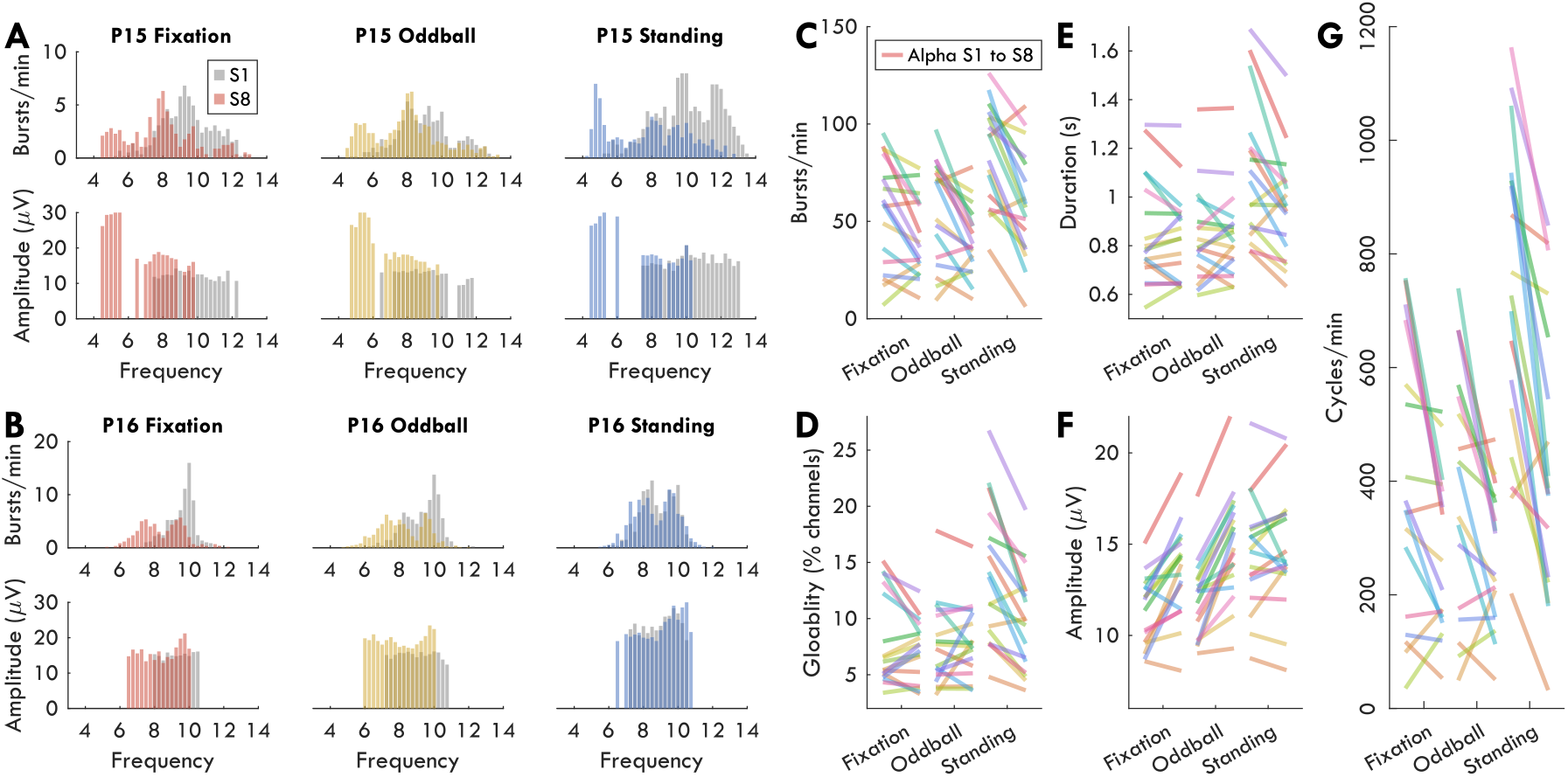
Distributions of bursts by condition. **A-B**) Distribution of number of bursts per minute for each frequency (top plot) and average amplitudes (bottom plot) for two participants (A, P15; B, P16). Gray histogram depicts data from the first extended wake recording (S1), and colored histograms the last (S8). Missing values in the amplitude plot correspond to bins for which there were fewer than 10 bursts across the 6 min recording. **C**) Alpha bursts per minute for all participants from S1 (left point of each colored line) to S8 (right point) for each condition. Each participant is a different color. **D**) Average alpha glob-ality, measured as the percentage of channels with an overlapping burst within ±1 Hz of the reference burst. **E**) Average alpha burst duration in seconds. **F**) Average alpha amplitudes, in microvolts. **G**) Average cycles per minute.

### Oscillation amplitudes increase with extended wake, but decrease during the WMZ

Figure 7 plots the change in theta (left column) and alpha bursts (right column) by average amplitude (top row), and cycles per minute (bottom row). Amplitudes tended to decrease after baseline sleep for theta (Fixation: t_(15)_ = −2.06, p = .057; Oddball: t_(16)_ = −1.99, p = .064; Standing: t_(16)_ = −2.55, p = .022). Amplitudes significantly decreased after sleep for alpha Fixation (t_(17)_ = −5.55), were trending for Oddball (t_(17)_ = −2.00, p = .061), and showed no change during Standing (t_(16)_ = 0.14). During extended wake, amplitudes increased substantially for both theta and alpha in the Fixation (theta t_(16)_ = 6.71; alpha t_(16)_ = 4.49) and Oddball conditions (theta t_(15)_ = 6.92; alpha t_(16)_ = 7.52). Theta amplitudes increased during wake in Standing (t_(16)_ = 2.15, p = .047) but no change was observed in alpha amplitudes during Standing (t_(16)_ = 0.17). The trajectory of the increase in amplitudes for both theta and alpha, and Fixation and Oddball, approximated that of an increasing saturating exponential function, with steeper increases at the beginning compared to end of the wake period. Against our expectations, however, both theta and alpha showed a robust decrease in amplitude during the WMZ for both Fixation (theta t_(16)_ = −3.19; alpha t_(16)_ = −3.71) and Oddball (theta t_(16)_ = −4.78; alpha t_(16)_ = −6.47).

**Figure 7:**
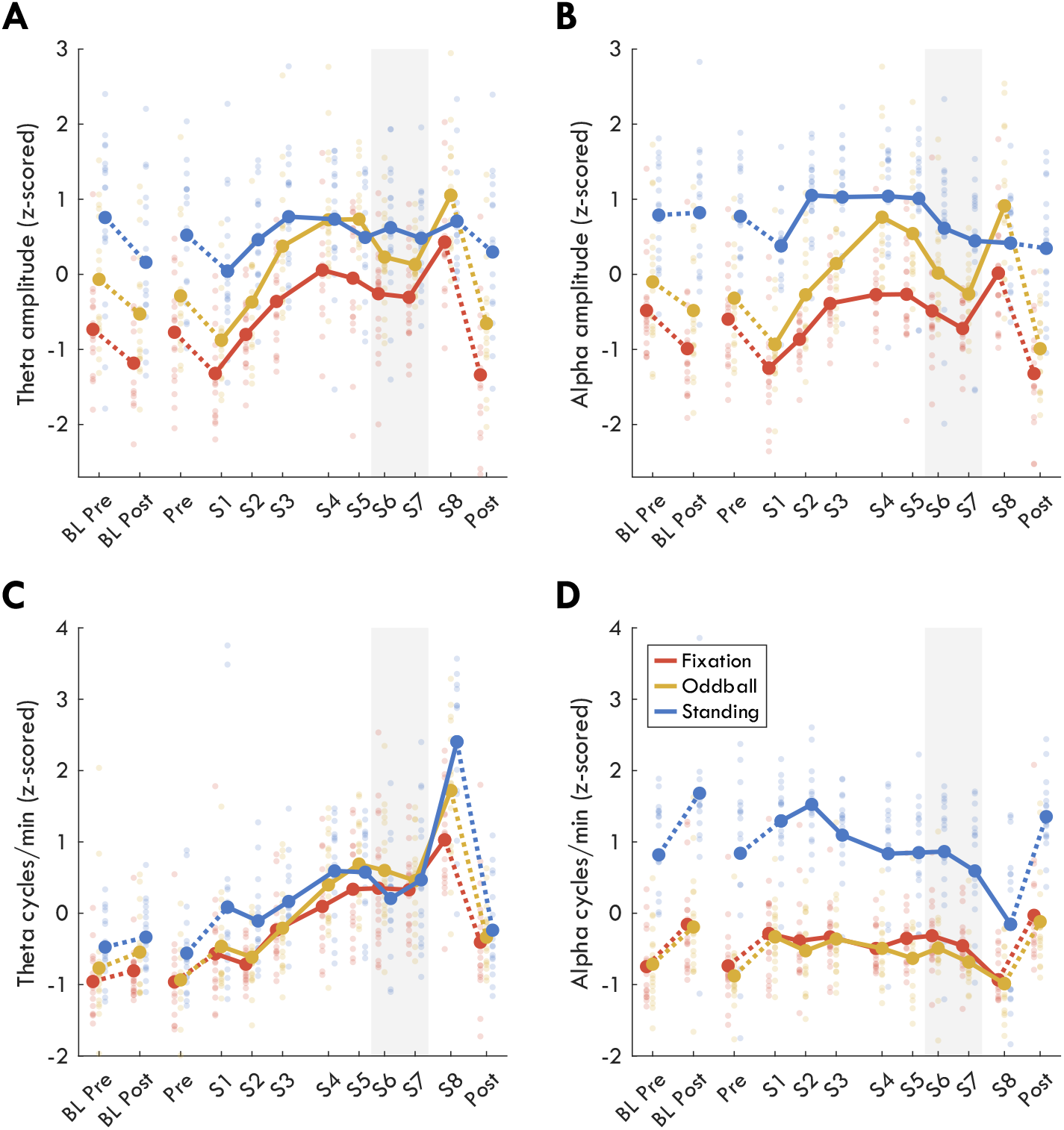
Z-scored burst changes in amplitude and quantity. **A**) Average theta burst amplitudes, **B**) alpha burst amplitudes, **C**) number of theta cycles per minute, **D**) alpha cycles per minute. Thick lines indicate group averages for each condition across sessions (x-axis), with color indicating condition. Solid lines connect sessions during the same-day extended wake period, and dashed lines indicate changes across sleep. S1-S8 are spaced out relative to the time they occurred within the 24 h wake period (Figure 1B). Dots reflect individual participants’ datapoints. The shaded gray area indicates the WMZ. All values are z-scored within each figure, such that sessions and conditions were pooled. Acronyms: BL, baseline; WMZ, wake maintenance zone.

Changes across baseline sleep in cycles per minute went in the opposite direction from amplitudes: theta quantities on average increased although this was not significant (Fixation t_(17)_ = 1.08; Oddball t_(17)_ = 1.25; Standing t_(16)_ = 0.67) and alpha significantly increased (Fixation t_(17)_ = 3.32; Oddball t_(17)_ = 2.63; Standing t_(16)_ = 3.63). During extended wake, theta quantities significantly increased in all conditions along a mostly linear trajectory (Fixation t_(16)_ = 5.57; Oddball t_(16)_ = 7.80; Standing t_(16)_ = 4.68), whereas alpha quantities decreased (Fixation t_(16)_ = −2.87; Oddball t_(16)_ = −3.31; Standing t_(16)_ = −6.17), primarily during S8. Theta and alpha cycles were significantly affected by the WMZ in opposite directions during the Standing (theta t_(16)_ = −5.02; alpha t_(16)_ = 2.51) and Oddball conditions (theta t_(16)_ = −4.03; alpha t_(16)_ = 2.91) but trending in Fixation (theta t_(16)_ = −1.77, p = .095; alpha t_(16)_ = 1.98, p = .065), such that theta quantities decreased relative to the overall trajectory, and alpha increased (or did not decrease along the expected trajectory).

To determine whether oscillation amplitudes and quantities originated from the same areas, we inspected the mean distribution of amplitudes and cycles per minute for theta and alpha bursts across the 123 channels, pooling sessions (Figure 8A-B). To determine whether the changes observed in Figure 7 were spatially dependent, we performed paired t-tests between S1 and S8 for each channel, with false-discovery rate (FDR) correction (Figure 8C-D).

**Figure 8:**
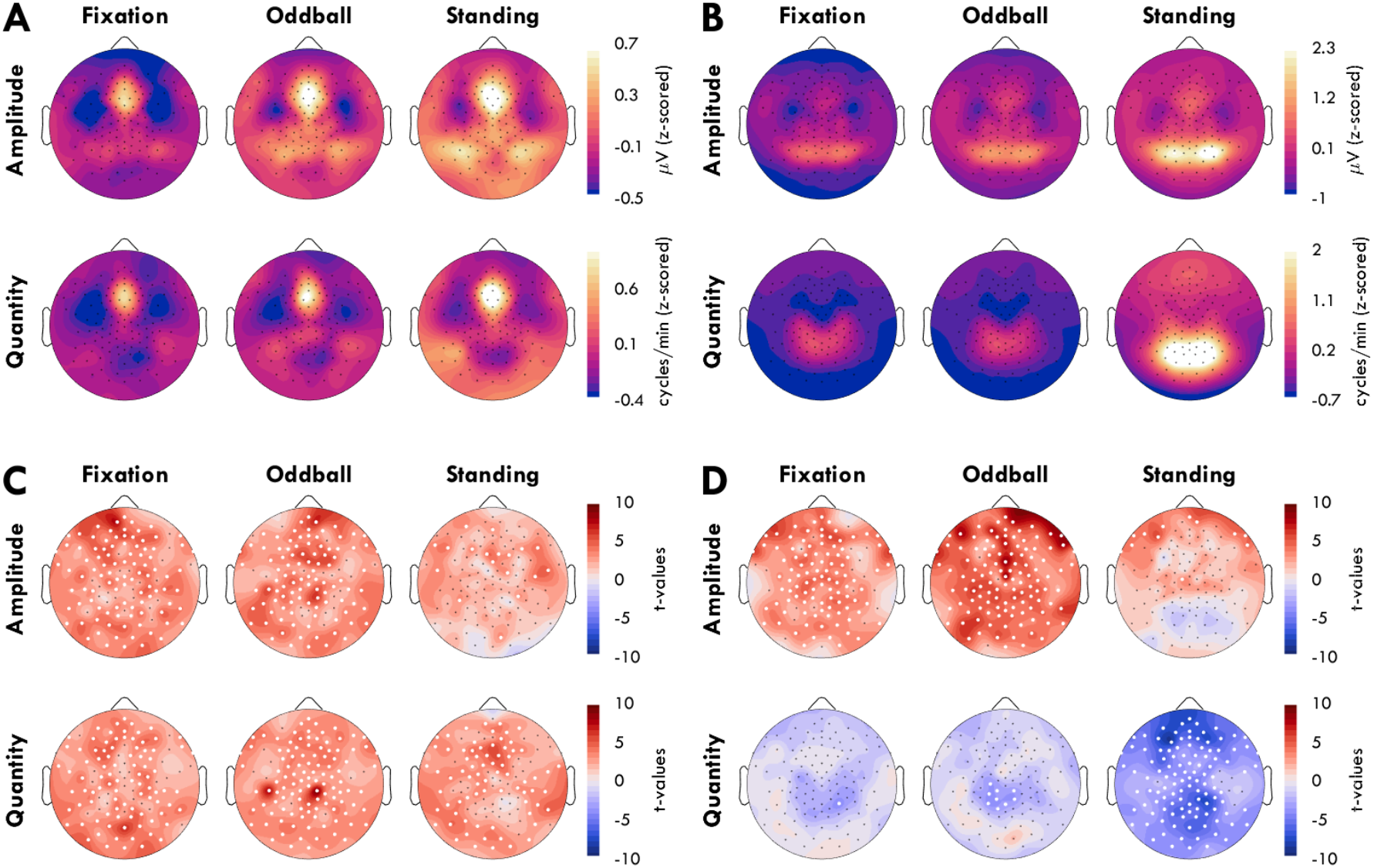
Topographic distribution of burst amplitudes and cycles per second. **A-B**) Amplitudes (top row) and cycles per minute (bottom row) for theta (A) and alpha bursts (B) across 123 channels for each condition, z-scored and averaged across all sessions. Warmer colors indicate higher amplitudes/quantities. **C-D**) Change in amplitudes and cycles/min from S1 to S8 for theta (C) and alpha bursts (D) represented as t-values, such that red indicates an increase with time awake. White dots indicate channels for which the difference was significant (p<.05, N = 17) based on paired t-tests, with false discovery rate correction.

For all three conditions, theta bursts were located primarily in frontal-midline channels, which also generated the largest amplitudes (Figure 8A). The increases observed with extended wake were wide-spread although somewhat patchy for both amplitudes and cycles per minute (Figure 8C), and were not limited to the main frontal-midline sources of theta. Alpha amplitudes and cycles per minute were instead spatially dissociated (Figure 8B), with high amplitudes originating more occipitally, and high quantities originating more centro-parietally. While the increase in alpha amplitudes in the Fixation and Oddball were similarly widespread as in theta, during the Standing condition the increase was only frontal (Figure 8D, top row). Instead, the decrease in alpha quantities was localized to the centro-parietal regions in Fixation and Oddball, with more widespread decreases in Standing (Figure 8D, bottom row).

### Mean pupil diameter and variance change during the wake maintenance zone, while pupil responses to oddball tones do not

To explore further what drives the different trajectories observed in the EEG data across wake, and in particular the changes in the WMZ, we analyzed pupillometry data from the Fixation and Oddball conditions. In both, mean pupil diameter after an initial decrease remained largely constant during the extended wake period (Figure 9A), with a specific increase during the two WMZ timepoints (Fixation t_(15)_ = 4.65; Oddball t_(12)_ = 2.17, p = .051). There was also a significant drop in pupil diameter from S1 to S8 during Fixation (t_(16)_ = −3.78, p = .002) but not Oddball (t_(12)_ = −1.39, p = .189). Interestingly, the two baseline recordings done in the evening an hour before bedtime (BL Pre, Pre) showed larger diameters during Oddball than during Fixation (BL Pre: t_(12)_ = 2.18, p = .050, g = 0.62; Pre: t_(16)_ = 3.22, p = .005, g = 0.63), but not during the same recording of extended wake (S7 t_(14)_ = −0.77, p = .453, g = − 0.16). This indicates an interaction, at least within the WMZ, between recording condition, pupil diameter, and prior sleep restriction.

**Figure 9:**
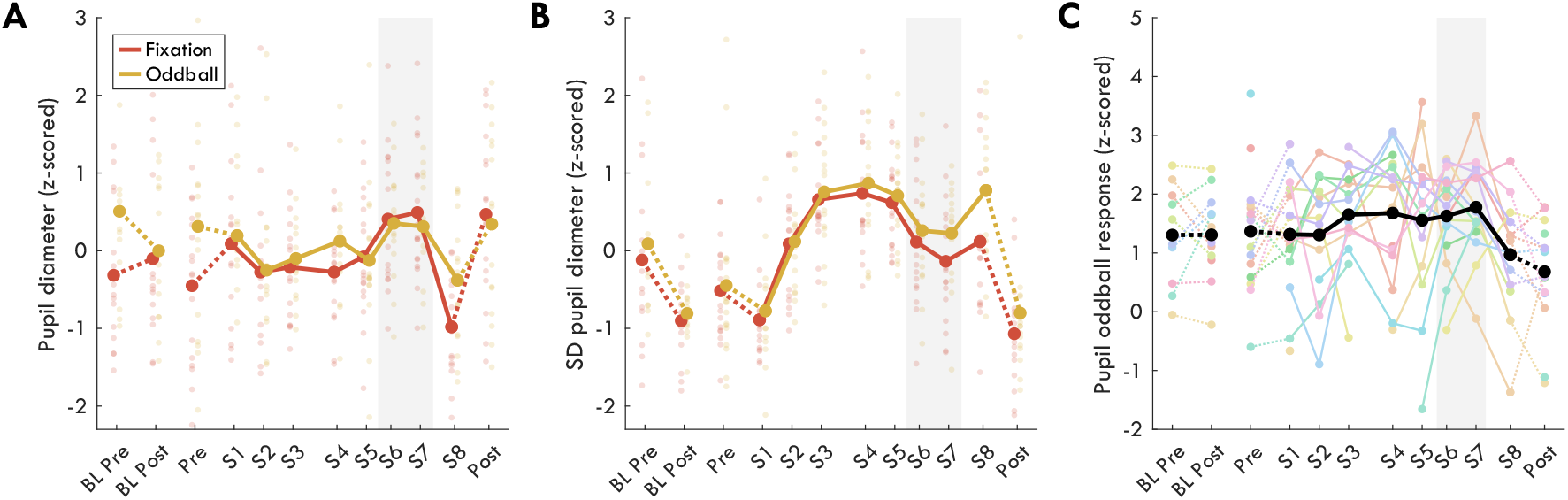
Pupil diameter. **A**) Mean diameter and **B**) standard deviations across the 6 minute recordings. Thick lines indicate group averages for each condition across sessions (x-axis). Solid lines connect sessions during the same-day extended wake period, and dashed lines indicate changes across sleep. Color indicates condition, dots reflect individual participants’ data-points. S1-S8 are spaced out relative to the time they occurred within the 24 h wake period (Figure 1B). The shaded gray area indicates the WMZ. All values are z-scored within each figure, such that sessions and conditions were pooled. **C**) Oddball pupil diameter response to target tones relative to standard tones from 0.5 to 2 s after tone onset. Colored lines indicate data from individual participants. Timecourses are provided in Figure 10. Acronyms: BL, baseline; SD, standard deviation; WMZ, wake maintenance zone.

Standard deviations of pupil diameter were also assessed (Figure 9B). Across sleep, there was a large significant decrease in standard deviations for both conditions (Fixation t_(13)_ = −3.76; Oddball t_(12)_ = − 3.66). During extended wake, standard deviations increased to maximum values in the afternoon (S4, 15:00). Standard deviations then tended to decrease during the WMZ, although the effect was only significant in the Oddball (t_(12)_ = −2.58, p = .024) and not Fixation (t_(15)_ = −1.69, p = .111). Across wake there was a significant increase in standard deviations from S1 to S8 (Fixation t_(16)_ = 3.52; Oddball t_(12)_ = 4.74), although S8 Oddball values returned to those of S3-S5 after the WMZ, whereas S8 Fixation remained lower.

Finally, we investigated the pupil response to target tones during the auditory Oddball condition (Figure 9C, Figure 10). Unfortunately, there was substantial data loss due to increased eye-closure with extended wake (Figure 11B) combined with equipment malfunctions during measurements, so power for this analysis is reduced. Pupil response to oddball tones was quantified as the area under the curve between the pupil response to targets relative to standard tones, from 0.5 to 2 s after tone onset.

**Figure 10:**
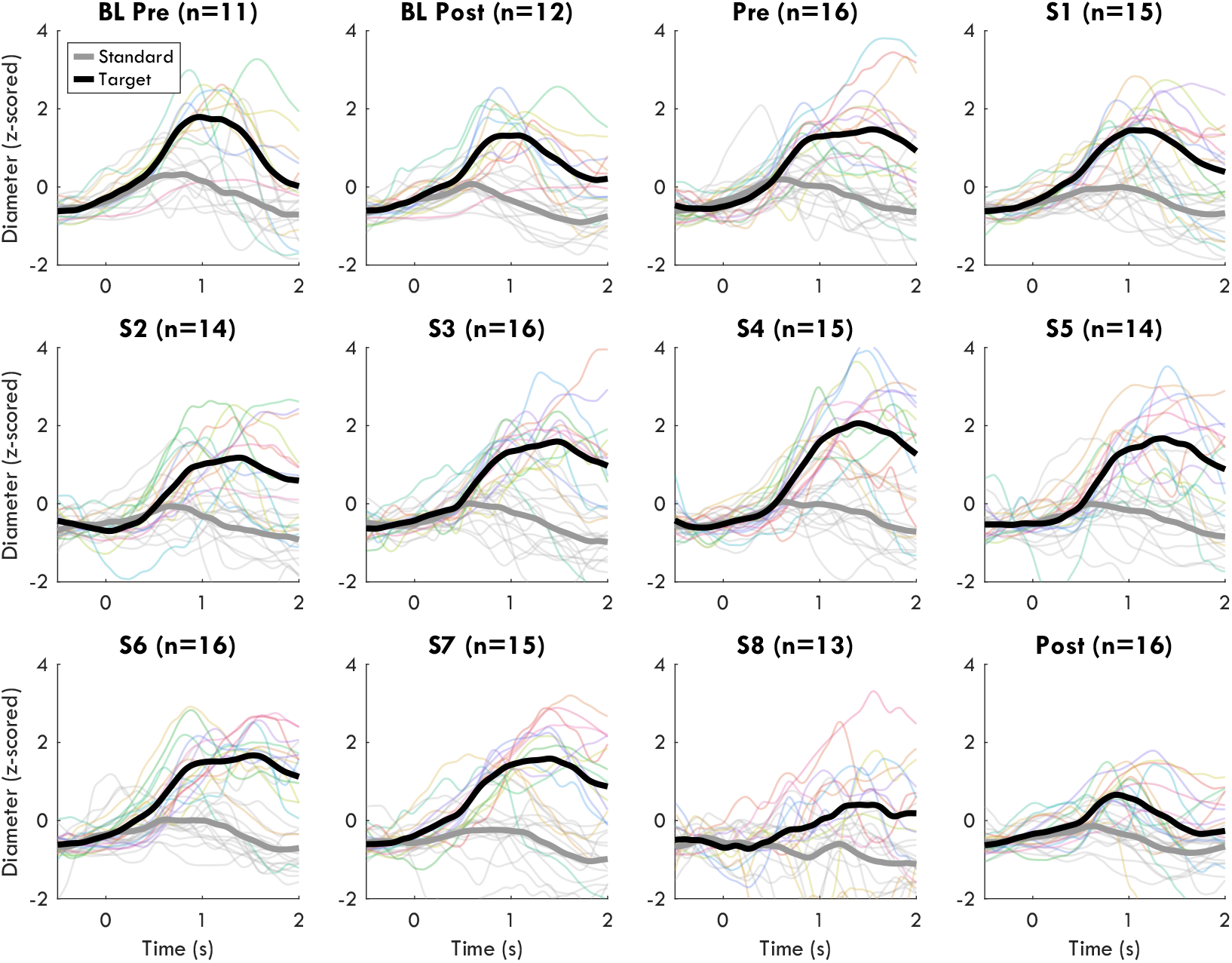
Pupil response to tones in the Oddball. Pupil diameters were locked to tone onset, baseline corrected (-.5 to 0 s from tone onset), and then z-scored pooling timepoints, tone type (target and standard), and session. Group average standards are in gray, oddball targets in black. Individuals’ response curves are thin gray lines, and individuals’ target responses are in thin colored lines. Timecourses were smoothed over 2 s for visualization. Due to data loss and increasing noise, multiple recording sessions were lost, and so the sample size for each session is indicated in the figure titles. Acronyms: BL, baseline.

**Figure 11:**
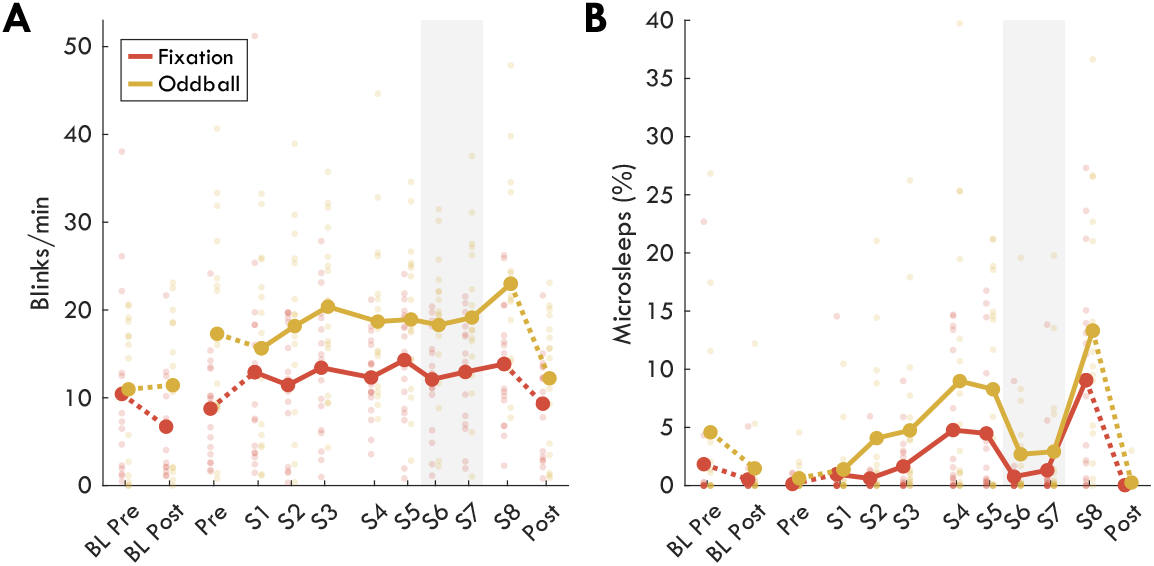
Eye-closures. **A**) Number of blinks per minute. A blink is defined as any eye-closure less than 1 s long. **B**) Percent of recording with ocular microsleeps (eyes closed longer than 1 s). N.B. here raw values are provided, although the statistics described in the text are with z-scored values. Thick lines indicate group averages for each condition across sessions (x-axis), with color indicating condition. Solid lines connect sessions during the same-day extended wake period, and dashed lines indicate changes across sleep. S1-S8 are spaced out relative to the time they occurred within the 24 h wake period (Figure 1B). Dots reflect individual participants’ datapoints. The shaded gray area indicates the WMZ. Acronyms: BL, baseline; WMZ, wake maintenance zone.

There was no change in response to targets from evening to morning around a baseline night of sleep (t_(9)_ = 0.26), nor was there a significant change from beginning to end of the extended wake period, although there was on average a decrease in pupil oddball response (t_(11)_ = −1.58). There was trending effect of the WMZ (t_(9)_ = 2.08, p = .067). However, as can be seen from Figure 9C, this was almost entirely due to a drop in the oddball response for S8 relative to previous sessions (e.g. S5 vs S8: t_(10)_ = 2.77, p = .020, g = −1.07). Furthermore, there was never a return to baseline values of pupil oddball response following recovery sleep (Pre vs Post: t_(13_) = 2.43, p = .030, g = −0.62), complicating any potential interpretation.

### Ocular microsleeps are sensitive to the WMZ, blink rates are not

In addition to actual pupil diameters, the same eye-tracking equipment could also be used to detect ocular behavior such as blinking and microsleeps, both of which increase with sleepiness (Crevits et al., 2003; Moller et al., 2006). We measured all eye-closures and split them into blinks when less than 1 s (Fatt & Weissman, 2013; Kwon et al., 2013), and as ocular microsleeps when longer than 1 s (Hertig-Godeschalk et al., 2020; Ong et al., 2013). We specify “ocular” to distinguish from proper microsleeps detected with EEG (Hertig-Godeschalk et al., 2020) and “behavioral” microsleeps detected as performance lapses (Poudel et al., 2014).

Blink rates (Figure 11A) gradually increased across wake in the Oddball (t_(13)_ = 4.37) but not Fixation condition (t_(16)_ = 0.62), and there was no change during the WMZ (Fixation t_(15)_ = −1.25; Oddball t_(15)_ = −0.21). On the other hand, the number of microsleeps (Figure 11B) went from near zero to 10-15% of the recording across extended wake (Fixation t_(16)_ = 4.81; Oddball t_(13_) = 4.18), and returned to <3% during the WMZ (Fixation t_(15)_ = −4.85; Oddball t_(15)_ = −6.28).

## DISCUSSION

The two-process model of sleep postulates that sleep need accumulates along an increasing saturating exponential function with time spent awake (Figure 1A), and the sleep homeostasis hypothesis proposes that this process is driven by increasing synaptic strength, which results in increased neuronal synchrony (Borbély, 1982; Tononi & Cirelli, 2003). Consequently, wake EEG oscillation amplitudes should reflect homeostatic sleep pressure.

Our results reveal that indeed both theta and alpha amplitudes follow an increasing saturating exponential function across extended wake, following the trajectory of sleep homeostasis (Figure 7A-B). Furthermore, both theta and alpha amplitudes returned to baseline levels following recovery sleep, with trending decreases for theta and highly significant decreases for alpha around baseline sleep. These results therefore match the predictions of the two-process model and the synaptic homeostasis hypothesis. It is still possible that synaptic strength does not drive this increase in amplitudes, but at the very least, changes in wake oscillation amplitudes correspond to the changes observed in slow wave activity (Borbély, 1982; Dijk et al., 1987).

In particular, the fact that the number of alpha oscillations decreased at the same time as amplitudes increased, and the former was widespread and the latter was localized (Figure 8D), clearly indicates a dissociation between these homeostatic changes in amplitude from whatever process causes oscillations to occur in the first place. More subtly, while both theta amplitudes and quantities increased during extended wake, the trajectories were different, suggesting again distinct mechanisms. Previously, two landmark studies found that theta power in resting state EEG increased locally depending on prior daytime activity (Bernardi et al., 2015; Hung et al., 2013). Re-analysis of these datasets may reveal that this effect was specific to oscillation amplitudes, even in the alpha band, which would further support the interpretation that the local effects were driven by synaptic plasticity.

### Sensitivity of different conditions to sleep homeostasis

In our data, while the Oddball condition followed the Fixation trajectories, the Standing condition did not. Theta Standing amplitudes did not increase past S3 during the extended wake period, and alpha Standing amplitudes neither decreased following sleep, nor did they increase during extended wake past S2. Given the overall larger amplitudes in this condition, this may be at least partially due to a ceiling effect, or a peculiarity of eyes-closed alpha. The occipital channels producing the highest quantity of alpha oscillations (Figure 8B) were also the ones who showed no increase in alpha amplitudes (Figure 8D), while frontal channels still showed a significant increase. At the same time, given the eyes-closed condition, it’s also possible that participants were closer to “true sleep” during S8 Standing. This is supported by the complete change in spectrogram in the Standing Back ROI (Figure 4). Altogether, our results indicate that oscillation amplitudes tend to reflect sleep homeostasis, however this is not the only factor contributing to these amplitudes. In practice, this means that different recording conditions will be more or less sensitive to changes in sleep homeostasis.

#### Potential mechanisms behind the WMZ

##### Cortical desynchronization during the WMZ

What did not match our predictions at all were oscillatory amplitudes during the WMZ. In both Fixation and Oddball, theta and alpha oscillation amplitudes decreased during the WMZ, then returned to their previous trajectories during S8. The effect sizes of the WMZ were actually larger for amplitudes compared to the number of oscillations (Table 01), with Fixation WMZ quantities only trending. The fact that oscillation amplitudes decreased implies that whatever mechanism drives the WMZ, it results in overall desynchronized cortical activity. In practice, this means that oscillation amplitudes can usually be used as a marker for homeostatic sleep pressure, except during the hours before habitual sleep. We don’t consider this dip in amplitudes during the WMZ and the difference in Standing oscillation amplitudes to indicate that the models themselves are inaccurate, just that amplitudes of oscillations are no longer accurately reflecting the underlying neuronal synaptic strength and sleep homeostasis.

##### Timing and duration of the WMZ

It is noteworthy how the WMZ briefly interrupts both the linear increase in theta quantities and the saturating exponentials of theta and alpha amplitudes. This highlights how the WMZ is limited in time, unlike the gradual circadian component of the two-process model (Figure 1A). The model was established based on measures of alertness, core body temperature, and melatonin concentration, all of which changed approximately sinusoidally across 24 h (Åkerstedt et al., 1979). While none of our outcome measures were paralleling these sinusoidal fluctuations, almost all of them were clearly affected by the brief WMZ. We therefore suspect that a specific pathway is responsible for the WMZ, distinct from melatonin concentration and core body temperature, although still synchronized to the suprachiasmatic nucleus (SCN), the brain’s timekeeper (Aston-Jones et al., 2001).

The timing of the WMZ in our study diverges slightly from some previous studies which find the WMZ to occur 3-6 h before bedtime (McMahon et al., 2018; Shekleton et al., 2013; Zeeuw et al., 2018), but is in agreement with others that find the WMZ 1-4 h before bedtime (Dijk & Czeisler, 1995; Lavie, 1997). Studies like ours with later WMZ times were conducted under normal office lighting conditions (~150 lux, or no manipulation reported), whereas studies with earlier WMZ times were recorded under dim lighting conditions (<10 lux). As demonstrated by Gooley et al. (2011), brighter light shifts melatonin onset to around 2 hours later, likely explaining the difference in results. This therefore means that whatever light-induced mechanism delays melatonin onset, it also delays WMZ onset.

##### Neural pathways of the WMZ

Given that both the mean and standard deviation of pupil diameters were also strongly affected by the WMZ (Figure 9), it is likely that the ascending arousal system (AAS) is involved. The AAS includes the locus coeruleus (LC), the ventral tegmental area and substantia nigra, the dorsal and median raphe nuclei, and the basal forebrain (Lloyd et al., 2022). All these areas have been linked to changes in pupil diameter (Joshi & Gold, 2020; Lloyd et al., 2022; Reimer et al., 2016), and have widespread connections to the rest of the cortex. The LC in particular has been linked to transitory pupil responses such as the increases observed during oddball tasks (Aston-Jones & Cohen, 2005; Joshi et al., 2016; Murphy et al., 2014). Since we do not find an increase in pupil responses to oddball target tones in the WMZ (Figure 9C), this may suggest that of all the AAS, the LC is actually not involved in the WMZ. Alternatively, this may merely indicate that pupil oddball responses are not a reliable indicator of LC activity across time, and instead spontaneous pupil fluctuations such as those reflected in diameter standard deviations may be more representative. In this case, based on Figure 9B, LC activity may be subjected to the combination of homeostatic sleep pressure, circadian pressure, and the WMZ. Unfortunately, due to reduced power, these results are suggestive at best, and furthermore the link between LC and pupil diameters has not been unambiguously established. Hopefully future studies will be able to determine which nuclei are involved in the WMZ.

##### The role of the WMZ in humans

It may be difficult to investigate the intracortical mechanisms of the WMZ because it’s possible the WMZ does not exist in rodents. The WMZ may even be human-specific because we have long consolidated sleep, unlike most other species (Campbell & Tobler, 1984). By ensuring that individuals do not initiate sleep too early, the WMZ largely guarantees that the subsequent 8 hours of sleep are *completed* within the correct circadian window, thus maintaining continued synchronization with the overall circadian rhythm and environmental light-dark cycles. During normal wake, the WMZ may not be apparent or even necessary, however when homeostatic sleep pressure is unusually high (for example from insufficient sleep the night before), such a mechanism would be critical to maintain wakefulness until the onset of the correct sleep window. In polyphasic-sleep species such as mice and rats, the timing of sleep onset for any given sleep episode is less critical.

The WMZ needs to be investigated more. As speculated by Strogatz et al. (1987), such a mechanism may be behind sleep disorders such as insomnia; if the WMZ never “shuts off”, this will result in substantially delayed sleep onset; if it is not present at all, this could result in circadian desynchrony. Taking this one step further, control over the WMZ could improve general wellbeing; being able to selectively shut it off could help with jetlag. Alternatively, activating the WMZ during night shifts could improve performance in critical industries such as emergency medicine or airline pilots.

#### Recommendations, limitations, and outlook

Our 4/24 study design has proven highly effective at capturing the WMZ, with multiple objective and subjective measures showing highly significant effects. The standard approach of starting with 8 h of sleep and then extending wake for more than 40 h does not often find effects during the first WMZ (McMahon et al., 2021; Shekleton et al., 2013), likely because normal wake is too close to ceiling. Given how burdensome 40 h of sleep deprivation can be, we recommend the use of sleep restriction for future research of the WMZ in both healthy and patient populations.

In addition to theta power, we found that the disappearance of ocular microsleeps is one of the best objective markers of the WMZ (Figure 11B), not previously described in the literature. These are quite easy to measure, requiring only eye-facing cameras. At the same time, while less sensitive, mean pupil diameter may instead be one of the more *specific* markers of the WMZ, as it was otherwise not as affected by increasing homeostatic sleep pressure, especially in the Oddball (Figure 9A). Other experiments have been conducted measuring pupil diameter during sleep deprivation, but did not find these effects in the WMZ (Daguet et al., 2019; Van Egroo et al., 2019). However, these studies measured only the first WMZ under well-rested conditions and the sleep deprivation was never sufficiently long to capture the second. Therefore, as with all other measures of the WMZ, it may be critical to record pupillometry under at least somewhat elevated levels of sleep pressure.

While we are satisfied with many of the design choices for this experiment, there is room for improvement. Given the unexpected importance of the WMZ, we would have benefitted from a traditional circadian marker such as melatonin concentration, which would have allowed more precise synchronization across participants to circadian phase. Additionally, more frequent recordings (e.g. every hour) would have provided a better temporal resolution to delineate the start and end of the WMZ for each participant. Regarding the wake recordings, given that the Oddball condition produced the largest effect sizes, its possible more controlled tasks provide better results than Fixation. Regarding the analyses, we implemented a relatively basic cycle-by-cycle burst detection algorithm, and there is ample room for improvement following more systematic development of this approach.

Regarding the interpretation of the EEG results, it is important that they are replicated with datasets using much longer bouts of sleep deprivation, spanning more than a single 24 h period and with more than a single recording after the WMZ. Likewise, data exists using the forced desynchrony protocol (Cajochen et al., 2002) which can properly dissociate circadian from homeostatic changes, as well as a constant routine protocol to control for homeostatic pressure (Cajochen et al., 2001). It would be important to see to what further extent amplitudes and number of bursts differently reflect circadian and homeostatic changes. Regarding the pupillometry results, it is important that they are replicated with larger sample sizes, and possibly comparing circadian changes with and without sleep restriction, as our results suggest an unexpected interaction.

#### Conclusions

In summary, by separately analyzing oscillation amplitudes and quantities, we were better able to validate the predictions of the two-process model of sleep and the synaptic homeostasis hypothesis. We demonstrated that both theta and alpha amplitudes increase with time awake, thus reflecting homeostatic sleep pressure. However, the wake maintenance zone proved to be such a potent contributor to wakefulness as to temporarily counteract these effects, impacting both the amplitudes and number of occurrences of oscillations. In addition to the EEG, we have identified ocular microsleeps as especially sensitive, and mean pupil diameter as specifically sensitive to this window. This specificity strongly suggests that the WMZ is caused by a wakefulness driver distinct from the gradual sinusoidal 24 h circadian fluctuations in alertness. Finally, we speculate that the ascending arousal system may be crucially involved, and that the WMZ may be human-specific.

## METHODS

Data from the task blocks of this experiment have been previously published in Snipes et al. (2022) where the overall study design, EEG preprocessing, and power analyses were established. Here, previously reported methods will only be briefly described.

### Participants

18 participants completed the experiment. University student applicants were screened for good health, good sleep quality, and at least some sleep deprivation vulnerability. 19 participants were recruited, and one participant dropped out midway. Of the 18 participants who completed the experiment, 9 were female and 3 were left-handed. Mean age (± standard deviation) was 23 ± 1 years old. All self-reported no hearing impairments. Data collection and interaction with participants was conducted according to Swiss law (Ordinance on Human Research with the Exception of Clinical Trials) and the principles of the Declaration of Helsinki, with Zurich cantonal ethics approval BASEC-Nr. 2019-01193.

### Experiment Design

Participants conducted a 4/24 extended wake paradigm, depicted in Figure 1. This involved habituation to a regular sleep-wake schedule prior to the experiment (minimum 4 nights), with bedtimes and wakeup times selected to match the participant’s preferred window and daily schedule. During the experiment, participants went to bed at their habitual bedtime, and were woken up 4 hours later. They were then kept awake for 24 h, followed by a recovery night. In addition to the extended wake bout, participants conducted a baseline night in which they slept during their habitual sleep window. The baseline was conducted before the extended wake bout in all but four participants.

The experiment schedule is in Figure 1B. Resting wake EEG recordings were measured before and after each night of sleep, and an additional 6 times during the 24 h wake period, for a total of 12 recordings. Prior to recordings S2-S7, participants were seated in the same position, watching 2 TV episodes around 40 minutes each, from a series of their choice. After each rest recording, participants were free to move around, and were provided with a home-cooked meal which they had selected from a list of vegetarian options (each meal during each break was the same). 6 of these breaks were included, each around 40 minutes (adjusting for delays in the schedule).

During the rest recordings, participants were seated in a comfortable armchair with a footrest (IKEA strandmon) in a well-lit room (~150 lux at eye level) and had to maintain fixation on a 20 cm red cross placed ~3 m from their head, ~30 cm below eye-level. The timing of the three rest recording conditions is depicted in Figure 1C. Each session began with a Fixation period, in which their only instructions were to maintain fixation on the cross and stay awake. This was immediately followed by an active auditory Oddball, where two types of tone were presented: standards (160 tones), and targets (40 tones). Participants had to press a button whenever a target tone occurred, while maintaining fixation and staying awake. Each tone lasted 60 ms, and for each participant the tone was either 660 Hz or 440 Hz for the targets, and vice versa for the standard tones. The interstimulus interval ranged randomly from 1.8 to 2.4 s, with a minimum of 3 standard tones between targets. After the Oddball, participants were provided a questionnaire to fill out, including the Karolinska Sleepiness Scale (Åkerstedt & Gillberg, 1990). Finally, participants stood up from the chair and moved to lean against the wall and had a Standing period with eyes closed. The purpose of this condition was to have a long recording with eyes-closed, the most typical condition for alpha activity (Kirschfeld, 2005), without participants falling asleep. Caldwell et al. (2000) previously found that there was no effect on alpha activity when comparing seated to standing recordings across sleep deprivation during eyes closed.

### EEG preprocessing & power analysis

EEG data was recorded at 1000 Hz, with 129 electrodes including the Cz reference, using EGI HydroCel Geodesic Sensor nets. Four electrodes were external to the net, and were positioned on the mastoids and under the chin for sleep scoring (sleep architecture is available in Snipes et al. [2022]). Two electrodes (126, 127) were located on the cheeks and excluded, leaving 123 channels for EEG data analysis after re-referencing to the average. Preprocessing and data analyses were done using EEGLAB (Delorme & Makeig, 2004) and custom MATLAB scripts. Data was downsampled to 250 Hz and filtered between 0.5-40 Hz. Major artifacts were identified visually, and physiological artifacts (eye movements, heartbeat, muscle activity) were removed with independent component analysis (ICA). Details are provided in Snipes et al. (2022).

Power was calculated as power spectral density using Welch’s method with 8 s windows, Hanning tapered, 75% overlap. Power values for each participant and each frequency were z-scored pooling across sessions, conditions, and channels. Z-scored values were then averaged across all channels excluding the outer-edge electrodes (Figure 3), or into 3 regions of interest (ROIs): Front, Center, and Back, marked in Suppl. Figure 5-1 of Snipes et al. (2022). When plotting spectrograms, a 1 Hz lowess filter was used to smooth the signal (Figure 4).

### EEG burst analysis

Burst detection was conducted with custom MATLAB scripts adapted from Cole and Voytek’s (2019) cycle-by-cycle analysis, originally coded in Python. The pipeline described below is also provided schematically in Figure 12.

**Figure 12:**
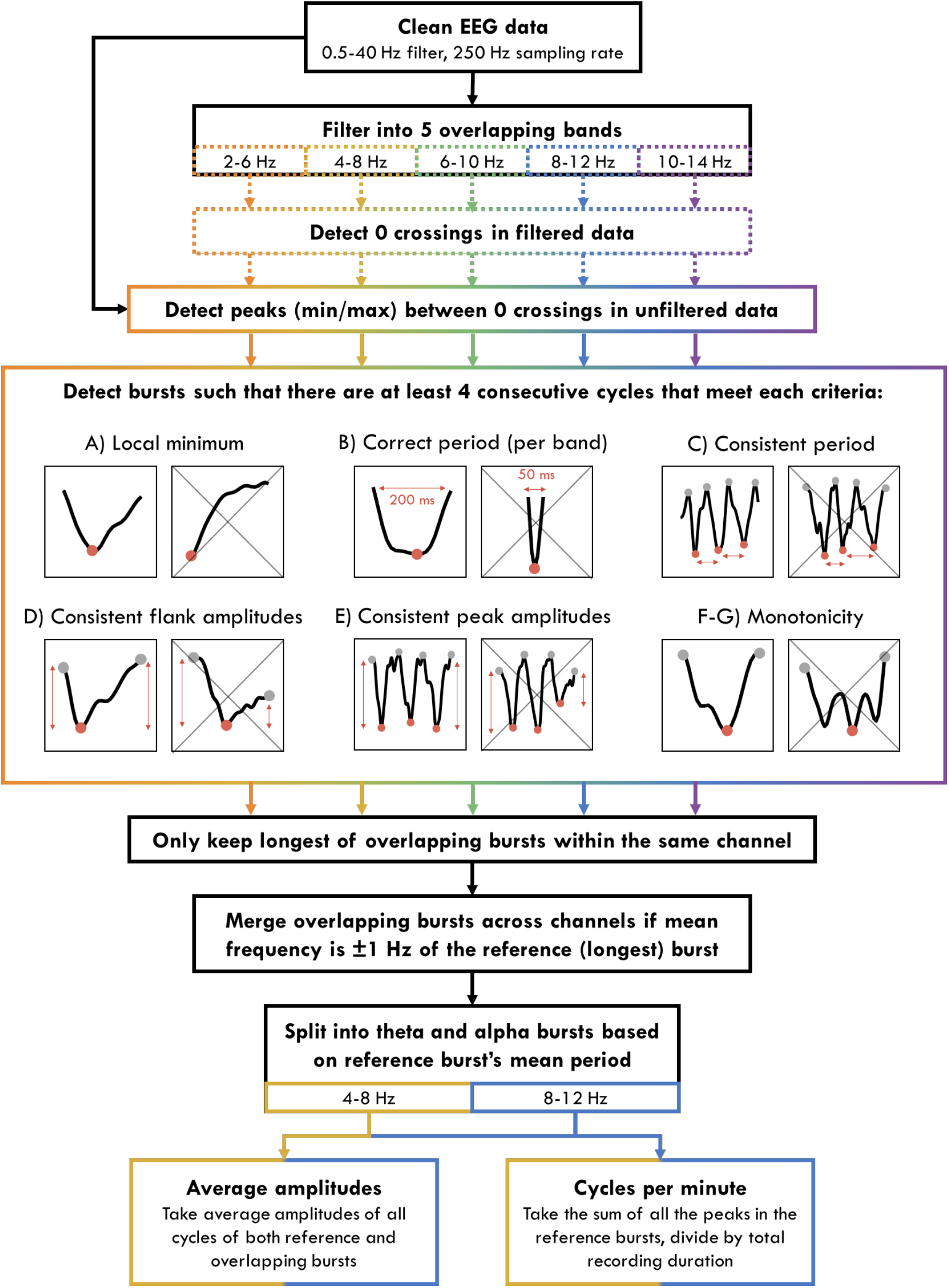
Burst detection algorithm. Solid outline indicates processes conducted on “unfiltered” data (only filtered between 0.5 and 40 Hz), dotted lines indicate processes conducted on filtered data (in 4 Hz bands). Colors indicate data processed separately for each band, black indicates processes done on pooled/undifferentiated data. A-G are examples of cycles that do or do not meet the criteria (crossed out). A and B depict half-cycles, from zero-crossing to zero-crossing. D and F/G indicate a whole cycle, from positive peak to positive peak. C and E indicate 3 consecutive cycles. In cycle-examples, red circles indicate the negative peaks, and gray circles positive peaks.

First, clean EEG data was filtered into narrow overlapping bands (2-6, 4-8, 6-10, 8-12, 10-14 Hz) using a minimum order high-pass then low-pass equiripple FIR filter (stopband frequency = passfrequency ± 1 Hz, passband ripple 0.04 dB, stopband attenuation 40 dB). Zero-crossings were identified in the narrow-band filtered data. Then between descending zero-crossings and rising zero-crossings, negative peaks were identified as the minimum value in the “unfiltered” data (minimally filtered during pre-processing between 0.5 and 40 Hz). Positive peaks were also identified as the maximum values in the unfiltered data between rising and descending zero-crossings, and these were used as the start and end of each cycle around the negative peak.

Once all the peaks were identified, 4 consecutive cycles had to meet the following properties in order to qualify as an oscillation burst: A) the cycle’s negative peak had to correspond to a local minimum; B) the mean distance to the neighboring peaks had to be within the range of the period of the filter (e.g. between 0.1 - 0.17 s when filtering between 6-10 Hz); C) the minimum ratio between the distance in time of the current peak to its neighbors had to be above 0.6 (i.e. similar consecutive periods); D) the rise amplitude, measured as the voltage difference between the prior positive peak to current negative peak, and decay amplitude of the cycle had to have a ratio of at least 0.5 (i.e. one flank was not less than half the amplitude of the other); E) the minimum ratio between the cycle amplitude (negative peak to positive peak voltage, averaging both neighboring positive peaks) and neighboring cycles had to be more than 0.6 (i.e. similar consecutive amplitudes); F) the proportion of timepoints decreasing in amplitude between previous positive peak and current negative peak, and timepoints increasing in amplitude between current peak and following positive peak, had to be above 0.6 (i.e. how much *time* during the cycle the signal went in the wrong direction); and G) the proportion of the voltage increasing from positive to negative peak, and decreasing from negative to positive peak, had to be above 0.6 (i.e. how much *amplitude* went in the wrong direction). The criteria B, E, and F are from Cole and Voytek, whereas A, C, D, and G are our additional optimizations. The parameters and burst-detection criteria were chosen through trial-and-error on an independent subset of data recorded during this experiment (the Game and PVT conditions of the SD task block reported in Snipes et al. [2022]). The procedure involved iteratively adjusting thresholds and introducing cycle exclusion criteria until the theta and alpha burst detection was largely consistent with visual inspection.

Bursts were detected for all frequency bands, using both the EEG signal and the negative of the EEG signal (because for mu-shaped rhythms, the sharper peaks resulted in better burst detection). Within each channel, overlapping bursts were compared, and the largest was retained intact. Smaller partially overlapping bursts were cut, and if the non-overlapping segment still retained 4 cycles, it was considered a new burst. Then, bursts were aggregated across channels based on temporal overlap (at least 50%) and if the mean frequency was within 1 Hz for the overlapping cycles.

Burst frequencies were defined as the reciprocal of the mean distances between negative peaks. Burst amplitudes were calculated by first averaging the rise and decay amplitudes of each cycle, then the average of these across all cycles in the aggregated bursts in different channels, then averaging the amplitude of all bursts within the band of interest. Burst quantities were calculated as the sum of all the cycles in the reference burst (the longest of all the overlapping bursts), divided by the duration of the recording, resulting in cycles per minute. This was chosen instead of the total number of bursts per minute because in extreme cases, bursts could become so long that their quantity decreased, and this was no longer representative of their occurrence in the data.

### Eye tracking & pupillometry

Eye tracking was done with Pupil Core “glasses” (Pupil Labs, Berlin, Germany). These were eyeglass frames with two rear-facing infra-red cameras. Pupil diameter was estimated from the video recorded with a sampling rate of on average 120 Hz. Data was exported using Pupil Player. All analyses were then conducted with a sampling rate of 50 Hz. During measurements, the eye tracker failed multiple times, resulting in substantial data loss. Sleep loss further resulted in noisier data (more eye-closure, less fixed gaze, half-closed eyelids).

The eye tracking variables blink rate and ocular microsleeps were measured using the confidence values of the pupil diameter estimates (from 0 to 1): when model confidence fell below 0.5, this was considered an eye-closure. This approach was chosen based on our observation of the video relative to the model confidence. Consecutive timepoints with confidence values over 0.5 that lasted less than 50 ms were still considered eyes-closed, and consecutive timepoints under 0.5 and less than 50 ms long were considered eyes open. The cutoff to split blinks and microsleeps was based on previous research identifying microsleeps as short as 1 s (Hertig-Godeschalk et al., 2020).

2D pupil diameter was estimated from the eye videos offline with Pupil Player, measured in pixels. Between the recordings, the eye tracking glasses were removed and readjusted. Therefore, in order to compare mean pupil diameter across sessions, the diameters in pixels had to be re-scaled. For every video, a frame was selected (192 x 192 pixels, 4.5 x 4.5 cm), and the eye’s iris diameter was measured in centimeters (when viewed at an angle, a disk becomes an ellipse, and the largest diameter of the ellipse is the diameter of the disk). By using the human mean iris diameter (12 mm), a conversion factor was calculated between pixels and millimeters, and this was applied to all 2D pupil diameter measurements:

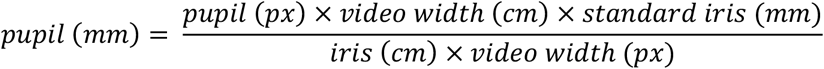

While this does not preserve individual differences in eye-size, it is sufficient for comparing across-session changes in diameter within participants (reasonably assuming irises do not change in size with sleep pressure). Furthermore, it allows the exclusion of unphysiological outliers.

Pupil preprocessing was done with the PhysioData toolbox (Kret & Sjak-Shie, 2019). Finally, removed datapoints less than 0.5 s were linearly interpolated, and then isolated chunks of datapoints less than 0.5 s were removed. Only data from one eye was used for each participant. The eye was chosen based on which had the most data after preprocessing.

To measure pupillary response to deviant tones during the Oddball, pupil diameters were epoched between −0.5 and 2 s relative to tone onset. All 40 targets were used, with 40 standards taken from the trial just prior to each target. Trials with less than 2/3 of clean timepoints were excluded. Recordings with less than 15 trials for either targets or standards were excluded. Furthermore, if any *timepoint* for a given tone type was derived by averaging fewer than 10 trials, this session was also excluded.

For each trial, the pupil response to tones was first baseline corrected (the mean between −0.5-0 s was subtracted from all datapoints in the trial), then all trials were averaged for each recording, split by target and standard tones. Participants with fewer than 6 recordings out of the 12 were excluded. Finally, average pupil responses for all timepoints, both targets and standards, and all sessions were z-scored within each participant. Pupillary response was calculated as the area under the curve between 0.5 and 2 s between target and standard.

### Statistics

All statistics were paired t-tests, with α = 5%. All t-tests were conducted on z-scored values (pooling sessions and conditions for each participant) so as to better account for interindividual differences, provide more normally distributed datapoints, and more fair comparison of effect sizes across outcome measures. Due to occasional data loss for different outcome measures, the degrees of freedom are always provided, from which the sample size can be inferred (N = DF+1).

Tests were selected a-priori for BL Pre vs BL Post to quantify overnight changes, and S1 vs S8 to quantify wake changes. To quantify the WMZ, a single value based on the average of the two recordings systematically showing effects (S6 and S7) were used. These were compared to an “expected” value based on S5 and S8, linearly interpolated. While previous studies quantified the effect by comparing WMZ values with measurements just prior, we considered this an under-estimate of the effect, as it doesn’t take into account the overall trajectory of the data, i.e. what values those timepoints would have had without the presence of the WMZ. However, our method can also overestimate the WMZ, if either S5 or S8 deviated substantially from the rest of the recordings. Therefore, results were interpreted in the context of the trajectories observed in the figures.

Hedge’s g effect sizes were reported for each test in Table 1, calculated with the Measures of Effect Size Toolbox (Hentschke & Stüttgen, 2011). Effect sizes are evaluated with Cohen’s rule-of-thumb (J. Cohen, 1988) such that g values <.2 are “small”, around 0.5 “medium”, and >.8 “large”.

No correction for multiple comparisons was done for these tests as the majority were rather exploratory (i.e. Oddball/Standing conditions), there were a mixture of dependent and independent comparisons, and the goal was comparing effects across outcome measurements (amplitudes vs quantities) rather than establishing significance. However, false-discovery rate correction (Benjamini & Hochberg, 1995) was conducted for the 123 t-tests in the topographies of Figure 8C-D.

## Code and data availability

Data used for Table 1 and Figures 2, 3, 7, 9 & 11 will be in Supplementary Materials, both untransformed and z-scored values. Additional unprocessed data is available upon request. All code is open source:

- The cycle-by-cycle analysis implemented in MATLAB: https://github.com/HuberSleep-Lab/Matcycle.
- The data analysis: https://github.com/snipeso/2Process_Bursts.
- The EEG preprocessing: https://github.com/snipeso/Theta-SD-vs-WM.
- The plotting graphics: https://github.com/snipeso/chART.
- The two-process model in Figure 1A: https://github.com/HuberSleepLab/2processmodel.
- The pupil preprocessing toolbox: https://github.com/ElioS-S/pupil-size

## ACKNOWLEDGEMENTS

This study was conducted as part of the SleepLoop Flagship project of Hochschulmedizin Zürich, with additional funding from the Swiss National Science Foundation (320030_179443) and Hirnstiftung. Marc Bächinger provided the eye-tracking equipment, technical assistance, and advice on the oddball protocol. Professor Christian Baumann and Niklas Schneider provided the EEG equipment. Simone Accascina helped with task programming and provided technical support during data collection. Noa Rieger aided with data collection. Lastly, a special thank you to all of our participants for taking part in this study.

## Notes

### Competing Interest Statement

The authors have declared no competing interest.

